# The topology of ecological drift

**DOI:** 10.64898/2026.01.12.698984

**Authors:** Ata Kalirad, Ralf J. Sommer

## Abstract

Ecological drift, the random fluctuation of abundance in communities, can drastically affect the possibility of coexistence. However, ecological drift has mostly been considered at the macro-ecological scale, whereas its potential contribution to short-term ecological dynamics remains underexplored. Here, we propose a stage-structured ecological graph theory (ssEcoGT) to include complex life cycles and multiple ecological interactions to study the topology of ecological drift. ssEcoGT enables rigorous exploration of how topology of a community can affect the possibility of coexistence between two species. Using ssEcoGT, it is possible to compare the behavior of a range of models – e.g., random graphs, random graphs with intraguild predation, and random graphs with rewiring – against a complete graph of the same size. Thus, ssEcoGT provides a new tool to explore the interplay between ecological drift, competition, and stage-specific interactions.

## Introduction

Ecological communities involve a finite number of organisms, inhabiting habitats that are shaped by variability in both space and time [1, 2, 3]. However, many of the often-used classical ecological models assume an idealized scenario of all-to-all interactions with little to no room for demographic stochasticity, i.e. random birth and death events, and their effects on the assembly of a community [4]. Ecological drift, the random fluctuations of abundance in finite communities, has received little attention, even though there exists an extensive theoretical body of work on coexistence [5].

The concept of ecological drift is generally associated with Hubbell’s unified theory of biodiversity [6, 7]. A drastic expansion of the theory of island biogeography [8], Hubbell suggests that much of macroecological patterns — e.g, species abundance distributions — could result from zero-sum ecological drift, that is, any dead individual is replaced by an offspring of any individual in the community, regardless of the identity of the individua [9]. In contrast to this *strong* interpretation, a *weak* interpretation defines ecological drift as a reflection of random birth and death events without necessitating an assumption of ecological equivalence of individuals [10]. When considering the combined effect of competition and drift on community dynamics – in contrast to macroecological patterns –, only the latter concept is adequate, since, depending on the nature of the interactions, e.g., competition over resource or intraguild predation, individuals in a community do not have equal chance of reproduction or death. Rather, the occurrence of these events results form a tug-of-war between competition and ecological drift [11, 12]. In this sense, the effect of ecological drift on the outcome of competition is analogous to the effect of genetic drift on the fixation of a beneficial mutation. The inescapable similarity between ecologcal and genetic drift, has led some authors to apply the tools developed for the latter to understand the former [13, 10]. However, this approach, in which effective population size becomes synonymous with community size, hinders the inclusion of various ecological interactions and complex community or metacommunity structures. A more ecology-oriented method is needed, given that spatial structure has been singled out as a key factor in shaping the ecological dynamics of a community [14, 15, 16].

Evolutionary graph theory (EGT) has convincingly demonstrated the importance of spatial structure in evolution [17]. Recently, we developed ecological graph theory (EcoGT), as an ecological equivalent to EGT, tailored to explore the possible coexistence in communities with static or dynamic spatial structure [18]. Here, we construct a stage-structured version of EcoGT, referred to as ssEcoGT. Crucially, ecological dynamics in ssEcoGT unfolds in a continuous rather than discrete fashion, which enables inclusion of complex life cycles and multiple ecological interactions.

### Stage-structured ecological graph theory (ssEcoGT)

In this study, firstly, we introduce the structure of the ssEcoGT model. Secondly, we systematically explore how community size, its structure, and various forms of competition, including intraguild predation, affect the strength of ecological drift.

Finally, we introduce an information-theory based quantification of the interplay between competition and drift for any given model, and argue for the need to emphasize a more rigorous distinction between census and effective community size.

### Components of ssEcoGT

Let us assume that graph G has been defined with a desired topology to simulate ssEcoGT. For the sake of simplicity, we assume that vertex *u_i_* designates a point in space and does not necessarily need to be occupied by an individual. Although this assumption does not ensure fixed population size, it enables the inclusion of stage-structured events as well as colonization in our model. Additionally, this assumption is necessary since the dynamics in ssEcoGT unfolds in a continuous as opposed to discrete fashion, e.g., EGT and EcoGT.

To illustrate how ssEcoGT works, we consider G to be a dodecahedron with 20 vertices and 20 edges. Vertex *v_i_* can be occupied by individual on vertex *u*, *x_u_* = (*g, d*), where *g* denotes its species identity and *d* denotes its developmental stage. For the sake of simplicity, let’s assume that an individual starts as a juvenile and develops into an adult. We start with 20 individuals of species *A* – all of which are juvenile – assigned to all *V* = {*v*_1_*, . . . , v*_20_} vertices. For individual *x_u_*, the only possible event is its development into an adult:

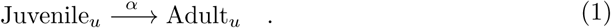

In our example population, with a spatial structure specified by dodecahedron G, there are 20 identical reactions representing development from juvenile to adult which can occur proportional to *α*. Gillespie’s algorithm [19] can be used to simulate this system of equations. Following the first-reaction method,

1. We draw *m* random numbers (*r*_1_*, … , r_m_*) for *m* = 20 reactions.
2. For reactions *i*, *τ_i_* is calculated:

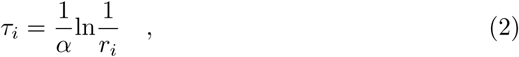

where *τ_i_* can be interpreted as the amount of time it would take until reaction *i* is executed.
3. The reaction with the smallest *τ_i_* is executed.
4. The simulation time *t* increases by the selected *τ_i_*.

Following the execution of reaction *r_u_*, *x_u_* is now an adult (Figure 1A). Following the execution of this reaction, our system includes 19 identical reactions that specify development from juvenile to adult, and a new reaction for *x_u_*:

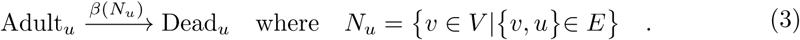

**Figure 1.**
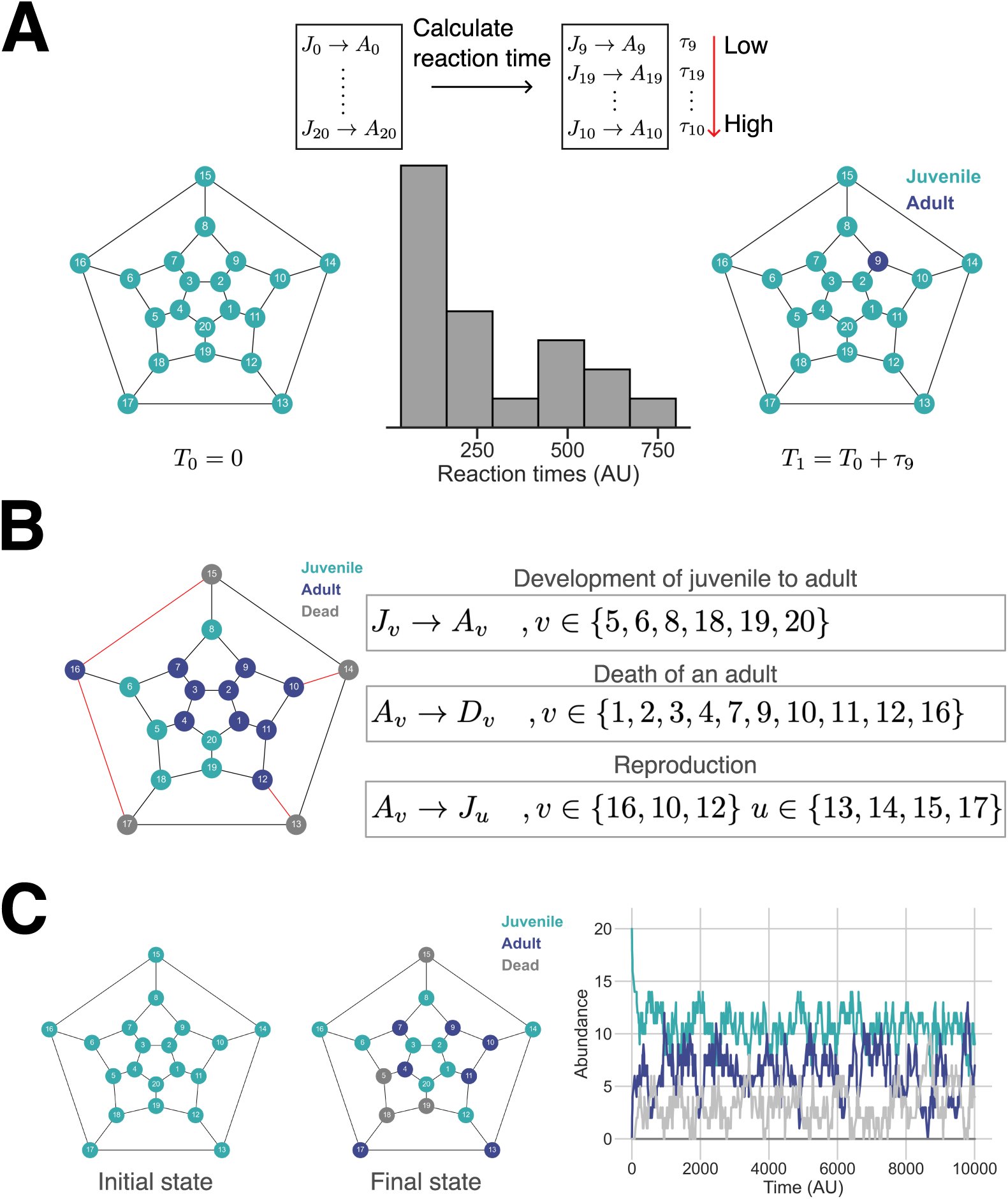
**(A)** In ssEcoGT, the possible events in a population are treated as reactions. In this case, at *T*_0_, all the vertices are occupied by juveniles and the only possible reaction is the development of a juvenile into adult. For a dodecahedron, for each of 20 juveniles on a graph, we assign a development reaction and calculate the reaction time for this reaction based on its propensity. The reaction with the lowest reaction time is executed, in this case reaction 9, resulting in an adult replacing juvenile on vertex 9. The current clock of the system is increased by the amount of time it took to executed reaction 9, *T*_1_ = *T*_0_ + *τ*_9_. **(B)** Given the current state of a vertex, different types of reactions can be assigned to it, where a vertex can have more than one reaction associated with it. In this example, three types of reactions are assigned to various vertices, with reproduction being possible only when an adult on vertex *v* is connected to an empty or, in this case, a vertex occupied by a dead individual. **(C)** A simulation of population dynamics on a dodecahedron, starting from an initial state where all the vertices are occupied by juveniles of the same species. For this simulation, the propensity of juvenile to adult reaction *α* = 0.003, the propensity of reproduction *f* = 0.02, and the parameter of connectivity-dependent death function *δ* = 0.005. The proportion of reactions and *τ* values for each reaction are shown in Fig. S1.

The rate of juvenile → adult reaction is a fixed value (*α*), whereas *β*(*N_u_*) is a function of the vertices connected to *u*, the vertex which is occupied by *x_u_*. In other words, development is independent of the identity of the neighboring individuals but death is. This assumption is not necessary – one can assume a connectivity-dependent development and a connectivity-independent death, amongst other possibilities – but it follows our previous EcoGT method, as well as the classic Lotka-Volterra species competition model, where death is dependent on interactions between species. The propensity of the death reaction is

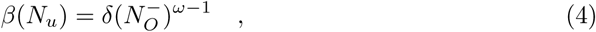

where *δ* is the baseline death rate 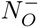 signifies the subset of vertices in *N_u_* that are occupied by allospecific individuals, and *ω* specifies the effect of allospecific neighbors on the death rate. This function differs from our original formulation of the death probability in EcoGT, since it scales the effect of interspecific interactions by the absolute number of allospecific neighbors – as opposed to the proportion of allospecific neighbors. Importantly, this formulation enables a more intuitive ecological interpretation of the intensity of competition as a function of competitors and is suitable when exploring the effect community size on its composition.

Reproduction is treated as a different type of reaction

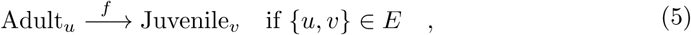

where an adult present in vertex *u* reproduces and creates a juvenile on vertex *v* with propensity *f* , provided *u* and *v* are connected ({*u, v*} ∈ *E*). For this reaction to be included in the set of all possible reactions on a given graph at the given time, there should exist an edge between vertices *v* and *u* and vertex *u* should either be empty or be occupied by a dead individual. Figure 1B illustrates the possible reactions given the state of individuals occupying the graph. A simulation of population dynamics in this model illustrates the emergence of steady-state distributions for each stage (Figure 1C).

Any additional stage-specific interaction can be added as a new type of reaction that is assigned to individuals. For example, consider a stage-specific predation scenario:

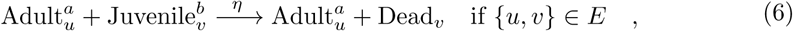

where an adult of species *a* on vertex *u* kills a juvenile of species *b* on vertex *v* with propensity *η*. Similar to reproduction, predation is conditional on the existence of an edge between a vertex occupied by a predator and one occupied by a potential prey. Figure 2A illustrate the connectivity-dependent predation in a scenario where two species exist in a community and both can prey upon allospecific juveniles.

**Figure 2.**
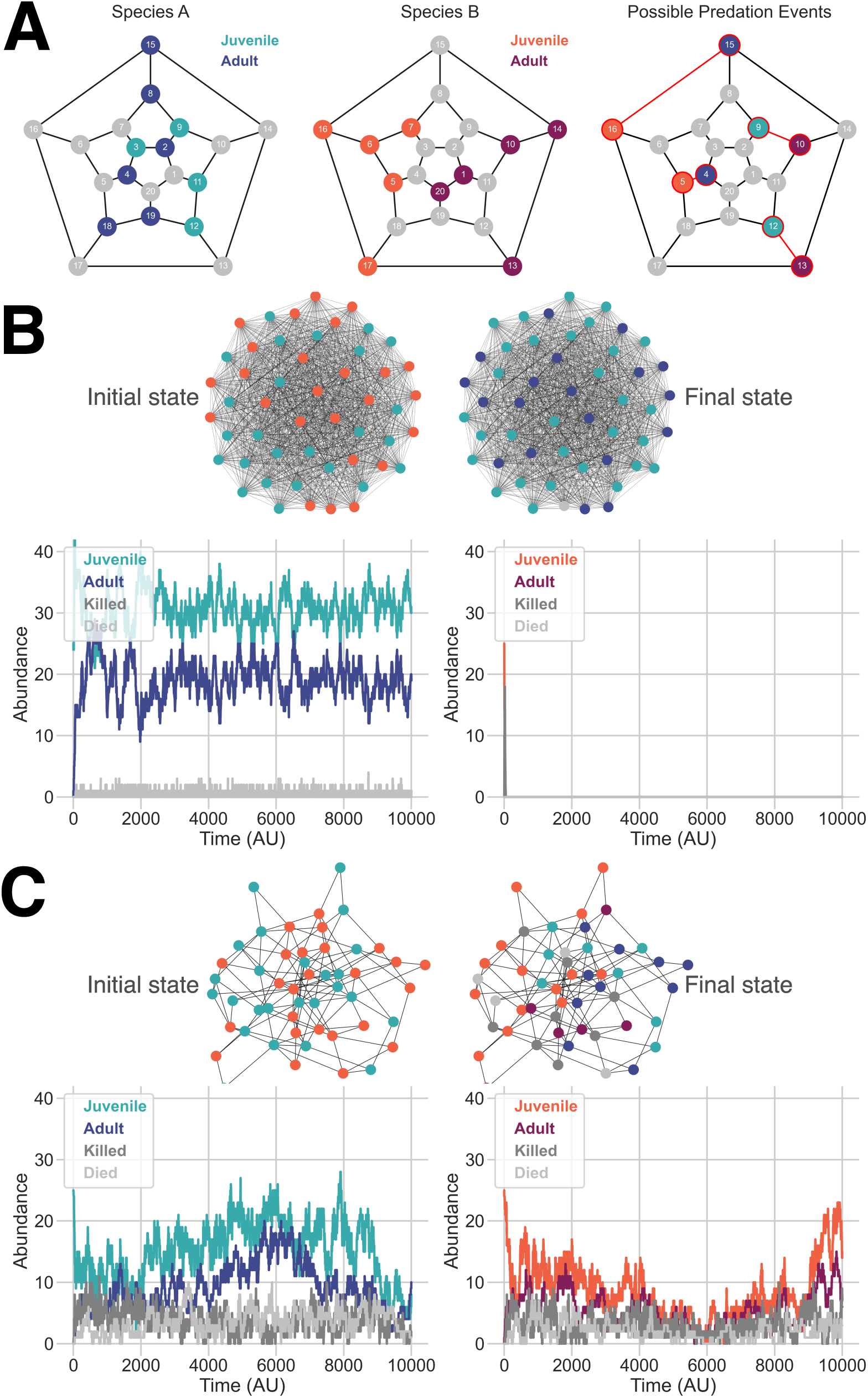
**(A)** An example of inclusion of predation in the stage-structured EcoGT. A community consisting of species A and B exists on graph 𝒢, a dodecahedron. Each species consists of juveniles and adults. In this scenario, we consider a situation where adults of species *i* can kill juveniles of species *j*, a case of reciprocal life-history intraguild predation (LH-IGP). In this community, 4 possible predation events exist, which can be simulated in ssEcoGT as four additional reactions. **(B)** A complete graph with 50 vertices is initialized with equal numbers of juveniles belonging to species A and B (initial state). In this scenario, the species interaction parameter *ω* = 0.5 for both species, meaning that the survival of an adult is positively influenced by the presence of allospecific neighbors. In the absence of reciprocal LH-IGP, this interaction parameter promotes coexistence, but when predation is included, ecological drift becomes possible, as either of the species can drive the competitor into extinction. In this realization, species A becomes monodominant (final state). **(C)** A community starting with 50 juveniles, belonging to species A and B in equal numbers, randomly assigned to graph 𝒢 which was created by the Erdőos–Rényi model with probability of edge creation between any two vertices *p* = 0.1. The absence of all-to-all interactions, in contrast to a complete graph, promotes coexistence of two species in spite of reciprocal LH-IGP. However, the observed coexistence could be transient. For **(B)** and **(C)**, the propensity of juvenile to adult reaction *α* = 0.003, the propensity of reproduction *f* = 0.02, and the parameter of connectivity-dependent death function *δ* = 0.005, and the propensity of predation reaction *η* = 0.05. For **(B)** and **(C)**, the proportion of reactions and *τ* values for each reaction during these simulations are shown in Fig. S2 and Fig. S3, respectively.

With the addition of predation, ssEcoGT allows for the simulation of complex scenarios in which the effect of resource competition – captured indirectly by including connectivity-influenced death events (Eq. 4) – can be distinguished from other forms of stage-specific interactions, for example, life-history intraguild predation (LH-IGP) [20]. As an illustration, consider a situation where the species competition, captured by *ω* in our model (Eq. 4) – analogous to parameter *α* in the classic LV model – is *<* 1. In the latter model, such parameter choice promotes coexistence. In EcoGT, a complete graph behaves in line with the classic LV model [18], since every vertex in a complete graph is connected to every other vertex, which mimics a well-mixed community. However, including reciprocal LH-IGP – where adults of both species can kill allospecific juveniles – in ssEcoGT results in ecological drift when two species compete on a complete graph, i.e., one of the two species becomes monodominant (Figure 2B).

Unsurprisingly, the topology of graph G on which a community resides greatly influences the interplay between competition and predation. This point can be illustrated by exploring the likelihood of coexistence of two species with *ω <* 1 and reciprocal LH-IGP on a random graph in which only a subset of possible edges are realized. For example, if G is created using the Erdőos–Rényi model, transient coexistence can be observed in spite of reciprocal LH-IGP (Figure 2C).

However, the influence of topology on the competitive outcome, specifically when including predation, hints at an important limitation, since predation is usually envisioned as an interaction coupled with exploration of the environment. In this respect, simulating competition alone could be justified on a community with a spatial structure that is fixed, but both predators and preys must be able to navigate the environment to provide a more realistic representation. We previously explored this idea by introducing rewiring algorithms in EcoGT [18]. In short, edges in a graph can be removed and added following algorithms that serve as abstract representations of realistic biological movements such as aggregation.

Since ssEcoGT is a continuous model, the rewiring can be also treated as a new type of reaction that occurs on a graph. In the simplest case, we can include random rewiring, which simply enables undirected movement of individuals. Consider G with *V* vertices. Assuming that all *V* vertices are occupied by living individuals – since empty vertices and those occupied by dead or killed individuals can not move – we can consider *V* reactions in our model in addition to existing reactions. For vertex *u*

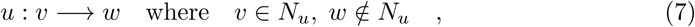

where *N_u_* indicates the set of vertices connected to *u*. Following this reaction, vertex *u* become disconnected from vertex *v* – randomly chosen from vertices connected to *u* – and is connected to *w*, randomly chosen from vertices that were not connected to *u*. It should be noted that vertex *w* could be occupied by dead or alive individuals or be empty. Given the nature of vertex occupancy in ssEcoGT, the number of rewiring reactions is dynamically changed as a function of the subset of vertices that are filled with living individuals.

Any type of rewiring algorithm can be included by modifying the way *w* is chosen when executing reaction 7. For example, if species prefer to aggregate, vertices occupied by conspecific individuals would be assigned a higher probability of being picked as *w*.

### Competition between species on a graph

Having illustrated the components of ssEcoGT, we now take advantage of this method to explore the effect of community size on the outcome of competition between species on a complete graph. The simplest scenario involves stage-structured competition between two species, without LH-IGP (Fig. 3A). Although all-to-all interaction is ensured in a complete graph, the finiteness of the maximum community size – the number of vertices (*V* ) – illustrates the effect of community size on ecological drift. In communities with *V* = 50 vertices, *ω_AB_* = *ω_BA_* = 1 always results in all-or-nothing outcomes, although few cases of coexistence are observed in *V* = 100, with coexistence becoming the expected outcome for this parameter combinations in larger communities (*V* = 200) (Fig. 3A). It would be tempting to conclude that community size serves as a predictor of ecological drift, similar to the relationship between the effective population size and the strength of genetic drift. In fact, community (or ecosystem) size has been used both in theoretical and experimental investigations of the effect of ecological drift on competition and extinction [21, 12, 22, 23]. However, the outcome of competition is expected to be influenced by both the number of vertices, which indicates the maximum size of the community, and the way those vertices are linked together, i.e., the structure of the community. Consequently, we suggest that the community size is analogous to the census population size, since both are correlate with the effectiveness of ecological or genetic drift, respectively. However, in order to provide an indicator of ecological drift akin to how effective population size predicts genetic drift, both community size and its topology should be taken into account.

**Figure 3.**
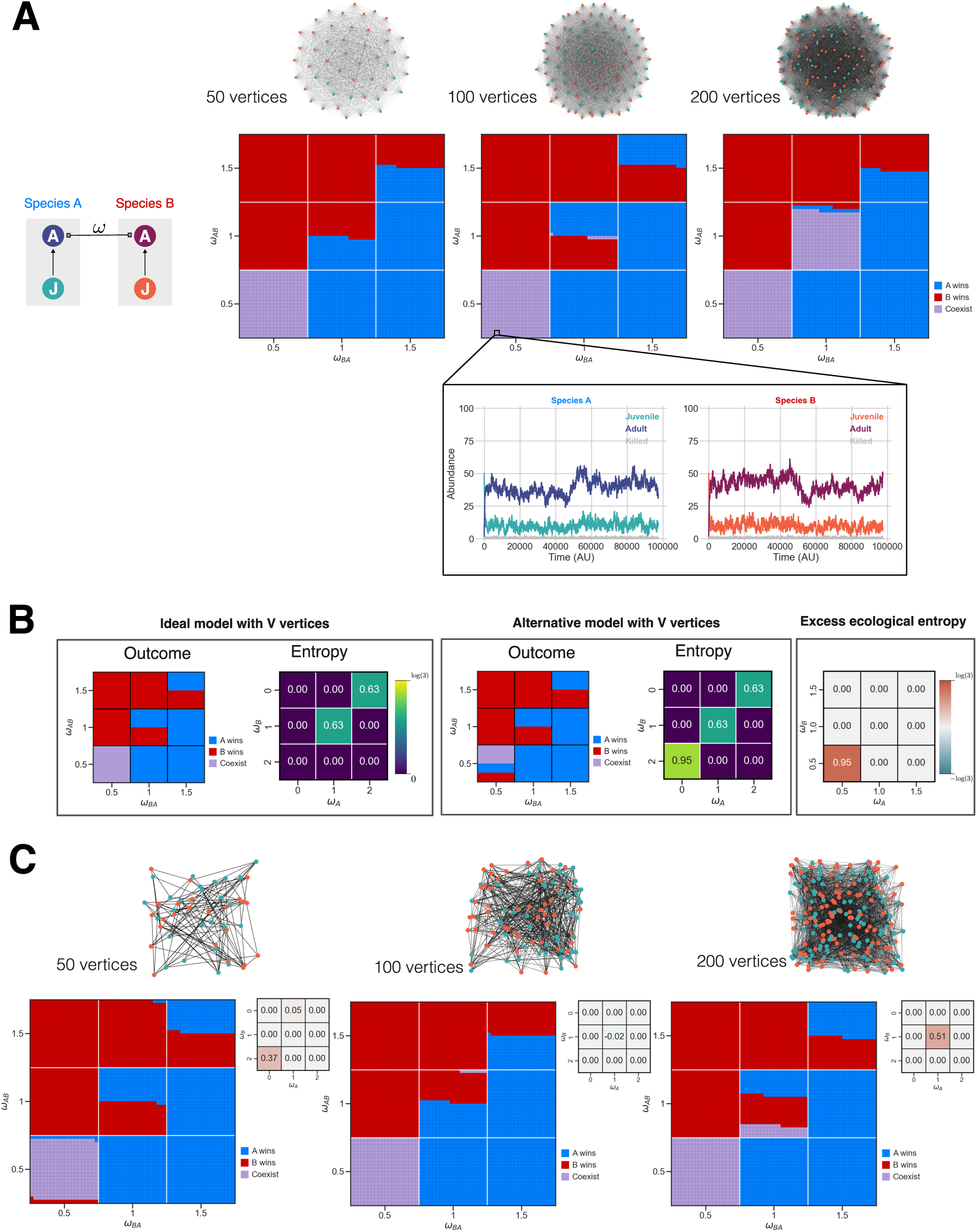
**(A)** The final community composition as a function of species competition parameters *ω_AB_* and *ω_BA_* without LH-IGP on a complete graph. Each pixel in this and subsequent plots indicates the final state of the community after reaching the maximum time 100000 (inset plot). **(B)** For any model, we can quantify its deviation from the expected outcome on a complete graph of the same size by measuring the entropy of simulations on a complete graph – as our ideal model – and the alternative model for a given *ω_AB_* and *ω_BA_* combination. The excess ecological entropy between the two indicates if the outcome of competition for the alternative model is more or less predictable compared to the ideal model. Since three outcomes are possible, the entropy is at most equal to log_2_(3). Normalized entropy was used to calculate the excess ecological entropy. **(C)** The results of competition on random graphs generated by the ER model with *p* = 0.1. For each graph size, the final state of the community after reaching the maximum time 100000 is juxtaposed next to the excess ecological entropy relative to a complete graph of the same size. Parameters used: *α* = 0.003, *f* = 0.02, *δ* = 0.005, and *η* = 0.05.

Much of our theoretical understanding of coexistence is derived from the classic LV model and its relatives. To illustrate the effect of community size and its structure, we suggest quantifying their effects by measuring the deviation of the outcome of competition of a given community with its size and its static or dynamic topology from the complete graph of the same size. One approach to provide such quantification is to borrow *excess entropy*, a concept commonly used in thermodynamics and information theory [24, 25].

In thermodynamics, excess property refers to the difference between an ideal gas and a non-ideal gas with respect to properties such as enthalpy, chemical potential, and entropy. For model *m* – i.e., a graph with a given number of vertices, defined topology, and the included interactions – we can calculate the entropy of the competitive outcome for a given combination of *ω_AB_* and *ω_BA_*

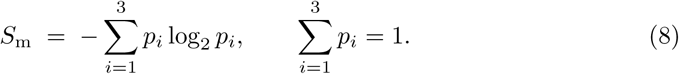

where 0 ≤ *S*_m_ ≤ log_2_(3). To quantify how different *m* is from our expectation, we compare model *m* of size *V* with a complete graph with the same number of vertices with only connectivity-dependent death for a given *ω_AB_* and *ω_BA_*. The complete graph serves as our ideal model since it mimics all-to-all interactions expected in the classic LV model. The difference between *m* and our ideal model is

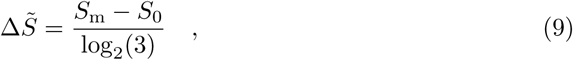

where *S*_0_ is the entropy of the ‘ideal’ model. This approach enables a simple visualization of the deviation of any model from a complete graph (Fig. 3B). Crucially, the entropy of a given model lends itself to an information-theoretical interpretation since it indicates the relative likelihood of three possible outcomes in a two species competition – i.e., species *A* or *B* winning or coexistence – and, in its excess entropy indicates how more or less predictable these outcomes are compared to a complete graph of the same size, the latter serving as an ecological equivalent to the ideal gas in the thermodynamic sense. Equipped with this quantification, we can compare the ecological outcome of competition in a wide-range of models with static or dynamic topologies with different type of ecological interactions. It should be noted that in contrast to excess entropy in thermodynamics, Δ*S̃* in. Eq. 9 can range between −1 and +1, since our ideal model does not have maximum entropy, in contrast to an ideal gas. Consequently, in order to avoid confusion, Δ*S̃* is referred to as *excess ecological entropy*.

Using the Erdoős–Rényi (ER) model, we can illustrate how topology and community size affect the community composition compared to a complete graph (Fig. 3C). For example, graphs with 50 vertices created by the ER model with *p* = 0.1 – resulting in the creation of ≈ 10% of the possible 1225 edges that exist in a complete graph with the same number of vertices – shows signs of ecological drift for *ω_AB_* = *ω_BA_* = 0.5, which is not observed in a complete graph of the same size (Fig. 3C). The smaller effective community size is more evident in graphs with 200 vertices, where a complete graph predicts *ω_AB_* = *ω_BA_* = 1 to be dominantly characterized by coexistence, whereas graphs of the same size generated by the ER model predict mostly all-or-nothing outcomes. In all these cases, excess ecological entropy clearly illustrates parameter combinations in which random graphs deviate from complete graphs (Fig. 3C).

Including reciprocal LH-IGP in communities on complete graphs promotes competitive exclusion, with the absence of coexistence in graphs with 50 vertices, given the combined effect of ecological drift and LH-IGP (Fig. 4A). This observation complicates the expected relationship between the community size and ecological drift, illustrating how complex ecological interactions would affect the effective community size, thus making a simple formulation of ecological drift – in the vain of genetic drift – ecological unrealistic at best.

**Figure 4.**
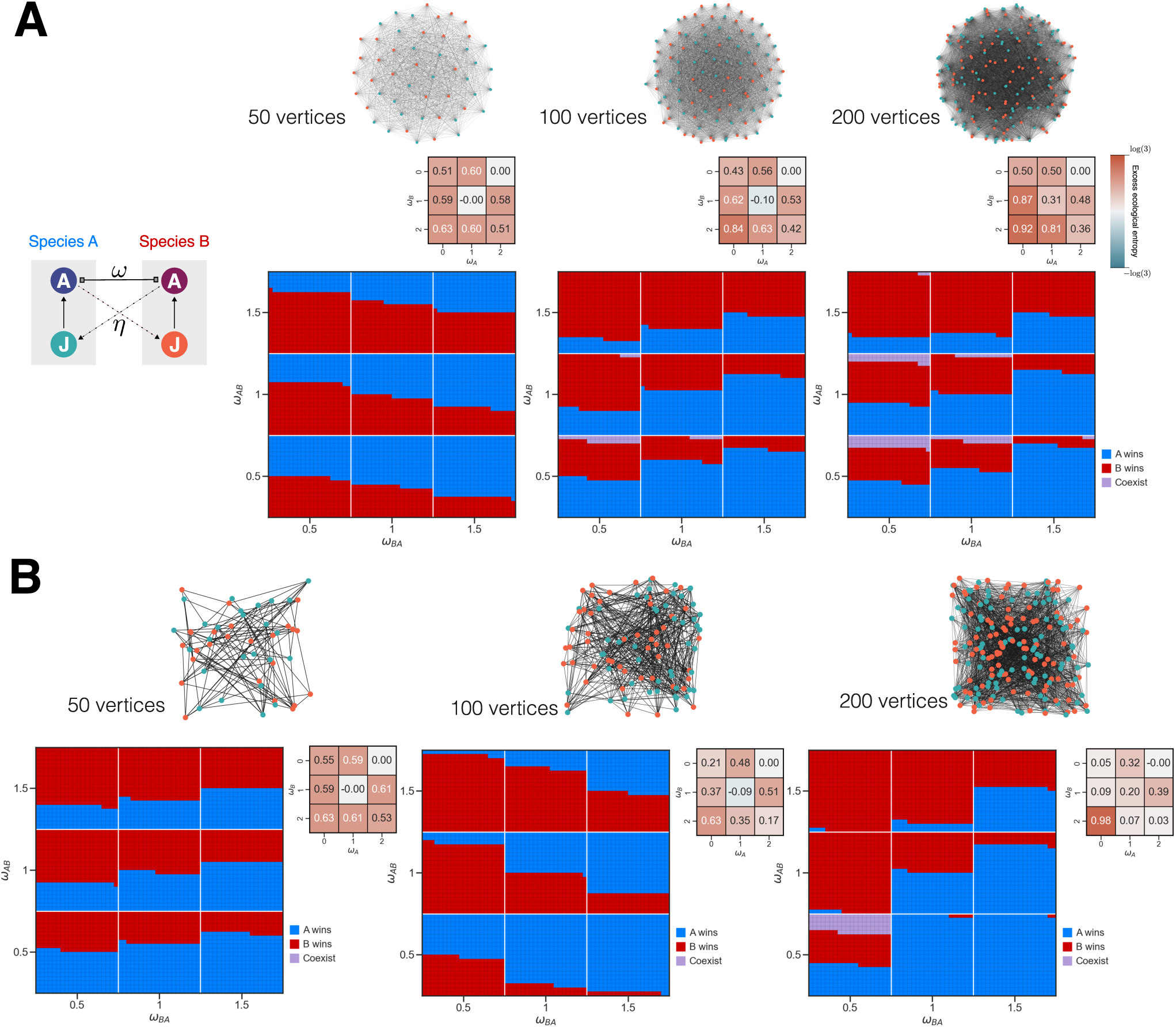
**(A)** The final community composition as a function of species competition parameters *ω_AB_* and *ω_BA_* with LH-IGP on a complete graph is juxtaposed next to the excess ecological entropy relative to a complete graph of the same size without LH-IGP. **(B)** The results of competition on random graphs generated by the ER model with *p* = 0.1 is juxtaposed next to the excess ecologcal entropy relative to a complete graph of the same size. Each pixel indicates the final state of the community after reaching the maximum time 100000. Parameters used: *α* = 0.003, *f* = 0.02, *δ* = 0.005, and *η* = 0.05.

The complexity comes into sharper relief when we turn our gaze on random graphs (Fig. 4B). Paradoxically, in random graphs with 200 vertices in *ω_AB_* = *ω_BA_* = 0.5 promotes coexistence more than the equivalent complete graph with reciprocal LH-IGP. This seemingly strange observation is due to the fact that the topology of a random graph, while reducing the effective community size, does reduce the possibility of predation as well. Consequently, while the ER topology promotes all-or-nothing outcomes with respect to competition, it alleviates the effect of predation.

## Conclusion

Ecological drift can greatly affect the outcome of ecological competition. However, despite a widespread recognition of its importance, a theory of “weak” ecological drift has been lacking. We have introduced a new model, ssEcoGT, as an extension of our previous EcoGT model, to enable rigorous exploration of how topology of a community – static or dynamic – can affect the possibility of coexistence between two species, quantified by the excess ecological entropy of a given graph with its specific set of interactions relative to a complete graph as an ideal point of reference. Using ssEcoGT, it is possible to compare the behavior of a range of models – e.g., random graphs, random graphs with LH-IGP, and random graphs with rewiring – against a complete graph of the same size (Fig. 5). Such a comparison illustrates the power of ssEcoGT as a tool to explore the interplay between ecological drift, competition, and stage-specific interactions such as LH-IGP.

**Figure 5.**
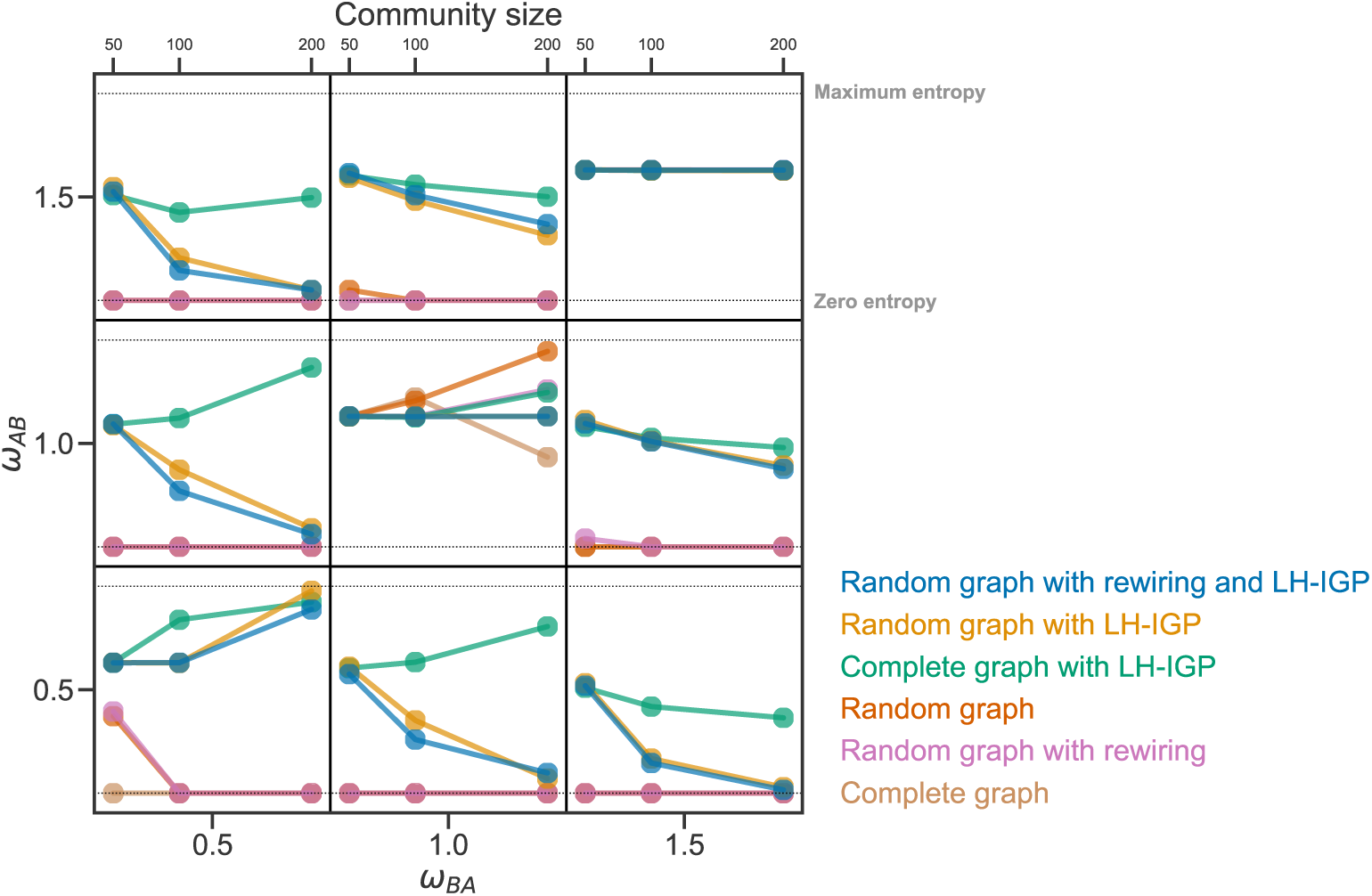
To illustrate the interplay between community size and community structure on the outcome of ecological competition, for each combination of *ω_AB_* and *ω_BA_*, we have plotted the entropy of the outcome of 400 simulations for each parameter combination for a given community size, ranging from 50 to 200. The complete graph serves as our point of reference, given its assumption of all-to-all interactions that approximates the classic LV model. The rewiring in the dynamic graphs are simulated as reactions that involve detachment of a random connected vertex from its neighbor and its connection to another random vertex in the community with propensity 0.01. Dotted lines are shown to indicate zero entropy – i.e., only a single outcome was observed for 400 simulations for a given graph with *V* vertices for a given *ω_AB_* and *ω_BA_* – and maximum entropy, where competitive exclusion of either of the two species and their coexistence is equally likely. Parameters used: *α* = 0.003, *f* = 0.02, *δ* = 0.005, and when LH-IGP was included, *η* = 0.05. The simulations for random graph with rewiring with and without LH-IGP are shown in Fig. S4.

The importance of demographic stochasticity – which leads to ecological drift – has been extensively discussed (Reviewed in [26]). However, even interpretation of classic experiments such as a series of experiments on *Tribolium* [27, 28, 29, 30], which indicate ecological drift, has been described in competitive terms, e.g., [31]. The relative obscurity of ‘weak’ ecological drift as an explanation can partly be attributed to the prevalence of mechanistic models derived from Lotka-Volterra equations or consumer-resource models and partly to the ubiquitous usage of ecological drift in the context of Hubbell’s unified theory. In this context, ssEcoGT provides a tool specifically suited to explore how ‘weak’ ecological drift can interact with various forms of competition and complex life cycles at community and – by extending the framework presented here to hypergraphs – metacommuniy levels. In this context, ssEcoGT could supplement the current efforts to construct an “ubermodel” for metacommunity ecological, enabling the exploration of metacommunity assembly in complex scenarios.

Finally, it should be noted that ssEcoGT can be utilized in the future to understand the role of phenotypic plasticity in community and metacommunity assembly. The role of plasticity and coexistence remains an under-explored area of community ecology research [32], and much of the difficulty in including plasticity in an ecological model stems from the complexity of interactions affected by plastic response to the environment. For example, during development from juvenile to adult in the nematode *Pristionchus pacificus*, a genetic switch makes the decision to develop either a narrow mouth form or a wide mouth form, the latter enabling the adult to bite and kill the juveniles of closely and distantly-related nematodes [33]. Incorporating such cases of plasticity in a model poses two major obstacles: (a) mouth-form plasticity in *P. pacificus* would unfold at a different timescale than birth and death and (b) the resulting predation is stage specific. However, ssEcoGT can easily overcome both these obstacles given its flexibility, which indicates the immense potential of this method in addressing complex ecological questions in the context of assembly and coexistence.

Thus, ssEcoGT provides a new tool to explore the interplay between ecological drift, competition, and stage-specific interactions.

## Data, Materials, and Software Availability

The software required to simulate ssEcoGT was written in Python (version 3.13) with NumPy 2.2.5 [34] and NetworkX 3.4.2 [35]. The code and notebooks to generate the results presented in this paper will be available at https://github.com/Kalirad/ssEcoGT.

## Acknowledgments

This work was funded by the Max Planck Society. We used the BIO/FML compute cluster at the Max Planck Institute for Biology Tübingen. We thank Andre Noll at Compute Cluster unit for the advanced support.

## Author contributions

AK and RJS designed the project. AK wrote the software, generated the results and visualized them. AK and RJS wrote the manuscript.

