## Supplementary figures for "The topology of ecological drift"

Supplementary Information

### Supplementary figures

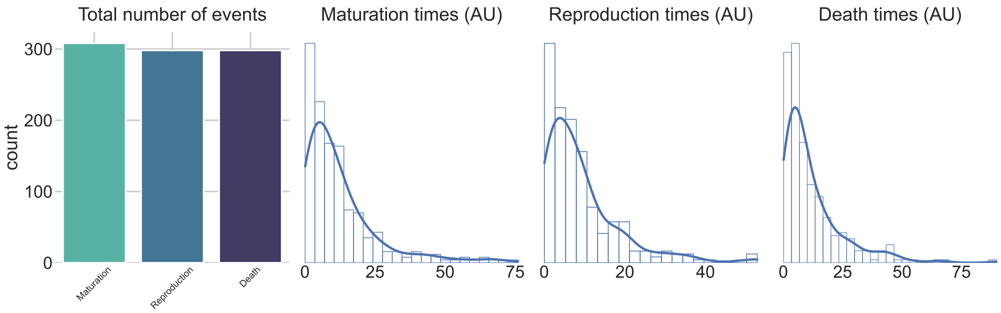


**Figure S1.** The final number of the three types of reactions executed during a simulation on a dodecahedron. The results are based on the simulation visualized in Fig. 1C. The histograms represent $\tau$ values generated for each type of reaction during the simulation.

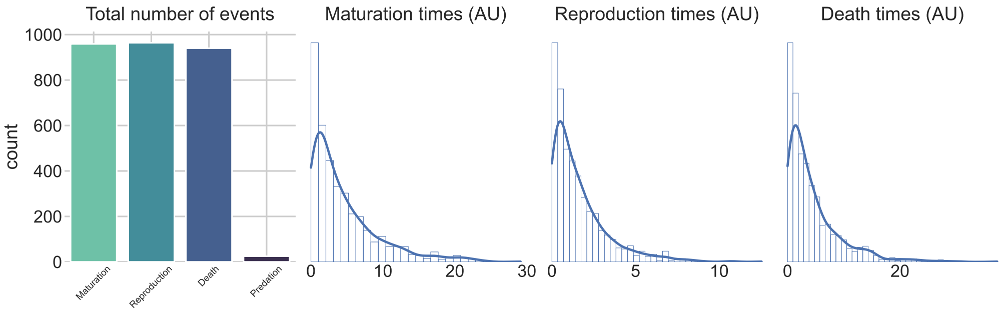


**Figure S2.** The final number of the four types of reactions executed during asimulation on a complete graph. The results are based on the simulation visualized in Fig. 2B. The histograms represent $\tau$ values generated for each type of reaction during the simulation.

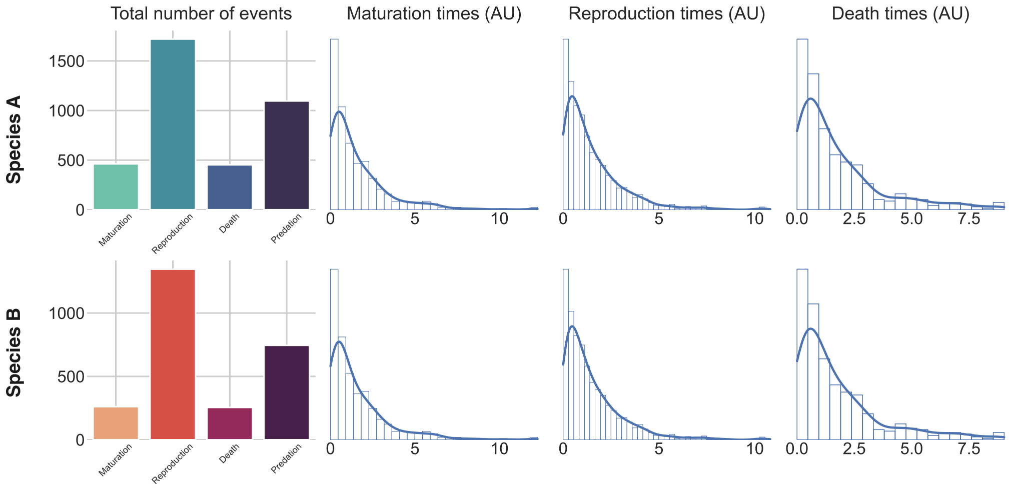


**Figure S3.** The final number of the four types of reactions executed during a simulation on a random graph created by the ER model. The results are based on the simulation visualized in Fig. 2C. Since in this simulation both species continued to coexist for the duration of the simulation, the number of reaction types and $\tau$ values are separately shown for each species. The predation reactions for species $i$ refers to the reactions in which an adult of species $i$ killed a juvenile of species $j$.

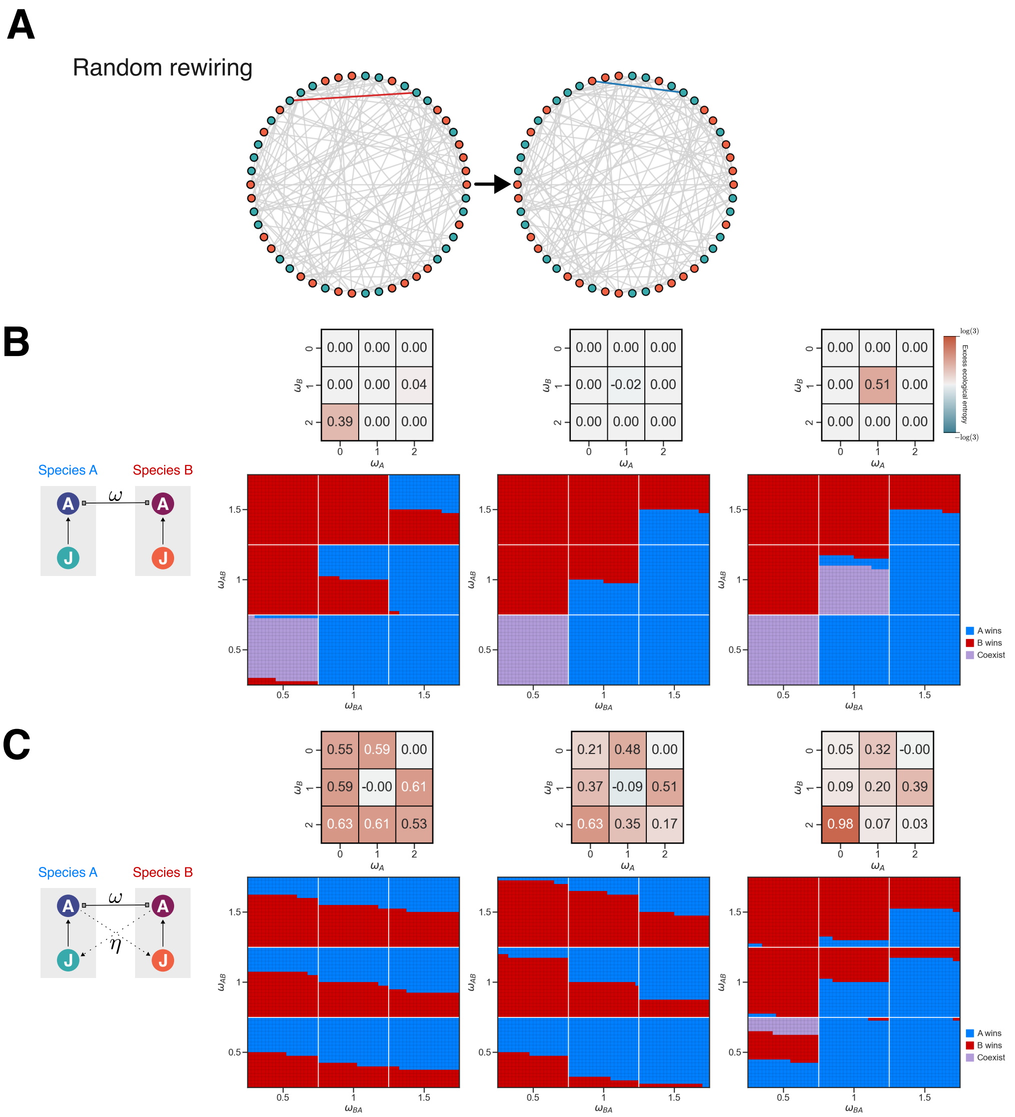


**Figure S4.** The interplay between rewiring of vertices on a graph and LH-IGP. **(A)** In random rewiring, a randomly-chosen vertex $u$, which is neither empty nor occupied by a dead individual, is disconnected from one of its neighbors and is connected to randomly-chosen vertex $w$ which was not connected to $u$. This algorithm preserves the total number of vertices in the graph during the simulation. (B) The final community composition as a function of species competition parameters $\omega_{AB}$ and $\omega_{BA}$ without LH-IGP on a random graph with rewiring is juxtaposed next to the excess ecological entropy relative to a complete graph of the same size. (**C**) The final community composition as a function of species competition parameters $\omega_{AB}$ and $\omega_{BA}$ with LH-IGP on a random graph with rewiring is juxtaposed next to the excess ecological entropy relative to a complete graph of the same size. The initial topology in each simulation was generated using the ER model with $p = 0.1$. Parameters used: $\alpha = 0.003$, $f = 0.02$, $\delta = 0.005$, $\eta = 0.05$, and rewiring propensity 0.01
